# Pretraining Enhances Megabase-Scale Gene Expression Prediction with GeneUnet

**DOI:** 10.64898/2026.08.13.744387

**Authors:** Ning Sun, William de Vazelhes, Pan Li, Tyler Katz, Jing Gong, Xingyi Cheng, Le Song, Eric P. Xing

## Abstract

Predicting gene expression from DNA sequence across diverse genomic tracks is essential for understanding gene regulation and interpreting non-coding variants. Existing supervised methods are limited to few species and fail to exploit conserved regulatory mechanisms, while DNA foundation models capture cross-species information but remain constrained to kilobase-scale contexts insufficient for this task. Here we introduce GB.GeneUnet, an 837M-parameter transformer-based U-Net pretrained on 6 trillion tokens from multi-species genomes in OpenGenome2, extending genomic context to 1 Mb with up to 100× inference speedup over GeneMoE, a preliminary MoE transformer baseline of similar model size pretrained on the same data. Fine-tuned for gene expression prediction, GB.GeneUnet achieves state-of-the-art performance on the Borzoi benchmark at 524 kb context, and attains performance comparable to AlphaGenome at 1 Mb context while requiring a lighter fine-tuning procedure. Together, these results establish a scalable framework linking multi-species pretraining to ultra-long-context gene expression modeling.

## Main

Sequence-based prediction of gene expression, which predicts functional genomic measurements across diverse biosamples at multiple resolutions directly from DNA sequence, is fundamental to deciphering the genetic regulatory code. Supervised models including Enformer (*1*), Borzoi (*2*) and the current state-of-the-art AlphaGenome (*3*) have achieved strong performance through joint training on human and mouse genomes. However, their scope is restricted to a small number of species and therefore underutilizes conserved regulatory principles preserved across evolution. DNA foundation models (*4–6*) provide a complementary paradigm. Self-supervised pretraining on multi-species genomes induces representations learning shared evolutionary constraints, enabling state-of-the-art performance on regulatory downstream tasks, such as variant effect prediction and promoter activity prediction (*5, 6*). These results suggest that cross-species representations could also benefit gene expression prediction. However, a key limitation remains. Accurate modeling of gene expression requires ultra-long genomic contexts exceeding 500 kb to capture distal regulatory elements, which surpasses the context capacity of existing transformer-based DNA foundation models. This limitation arises because standard transformer architectures scale quadratically with sequence length, making extension to megabase-scale inputs at billion-parameter regimes computationally prohibitive.

Here we introduce GB.GeneUnet, an 837M-parameter transformer-based U-Net designed to overcome the context length bottleneck inherent to existing genomic encoders. GB.GeneUnet adopts a hierarchical architecture that progressively downsamples sequence representations from 1 bp to 128 bp via stacked downsampling blocks and restores resolution through skip-connected upsampling blocks, enabling effective modeling of genomic contexts up to 1 Mb (**Fig 1.b**). GB.GeneUnet is pretrained on 6 trillion tokens from multi-species genomes in OpenGenome2 (*6*), then fine-tuned on two gene expression prediction benchmarks: the Borzoi benchmark (*2*) (524 kb context, 32 bp resolution) and the AlphaGenome benchmark (*3*) (1 Mb context, multi-resolution) (**Fig 1.a**). To isolate the architectural contribution of the U-Net design, we additionally pretrain GeneMoE—a preliminary MoE transformer encoder available in three model sizes (235M, 470M, and 1B parameters) supporting up to 256 kb context—on the same dataset as an architectural baseline (see Supplementary Sec. A)

**Figure 1.**
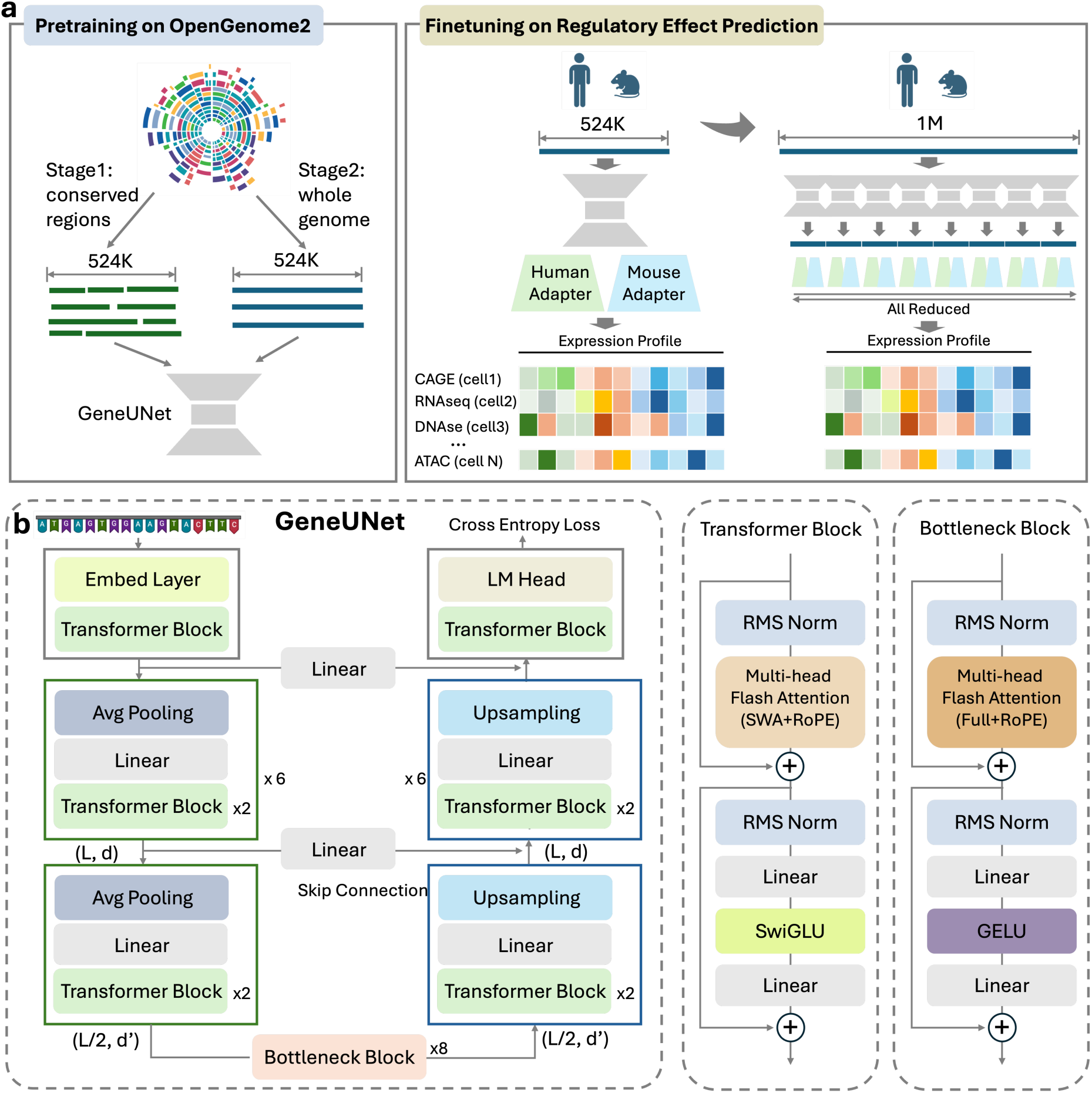
| a. Overview of the training pipeline. GB.GeneUnet is pretrained on multispecies genomic sequences from OpenGenome2 in two stages: an initial pretraining phase on conserved regions, followed by a midtraining phase extending to whole-genome data. The model is then fine-tuned for regulatory effect prediction using species-specific adapters in two phases, first at 524-kb context length and subsequently extended to 1-Mb context. **b. GB.GeneUnet architecture.** A Transformer-based U-Net with symmetric encoder–decoder structure, combining hierarchical downsampling, a global-attention bottleneck, and upsampling with skip connections to enable ultra-long-context modelling.

We first evaluate GB.GeneUnet against GeneMoE-235M and GeneMoE-1B on perplexity and inference efficiency across context lengths from 8 kb to 524 kb. GB.GeneUnet achieves consistently lower perplexity than both GeneMoE variants across all evaluated lengths, with the gap widening at 524 kb where GeneMoE exceeds its pretraining context limit (**Fig 2.b**). At matched token budgets, GB.GeneUnet also delivers substantially higher inference throughput: despite having 3.6× more parameters than GeneMoE-235M, GB.GeneUnet achieves 35× faster inference; against GeneMoE-1B, GB.GeneUnet achieves a 100× speedup while using only 84% of its parameters (**Fig 2.a**). This efficiency gain stems from the hierarchical compression of the U-Net architecture, which progressively reduces effective sequence length at deeper layers—in contrast to GeneMoE, which applies full-length attention at every layer despite sparse expert routing.

**Figure 2.**
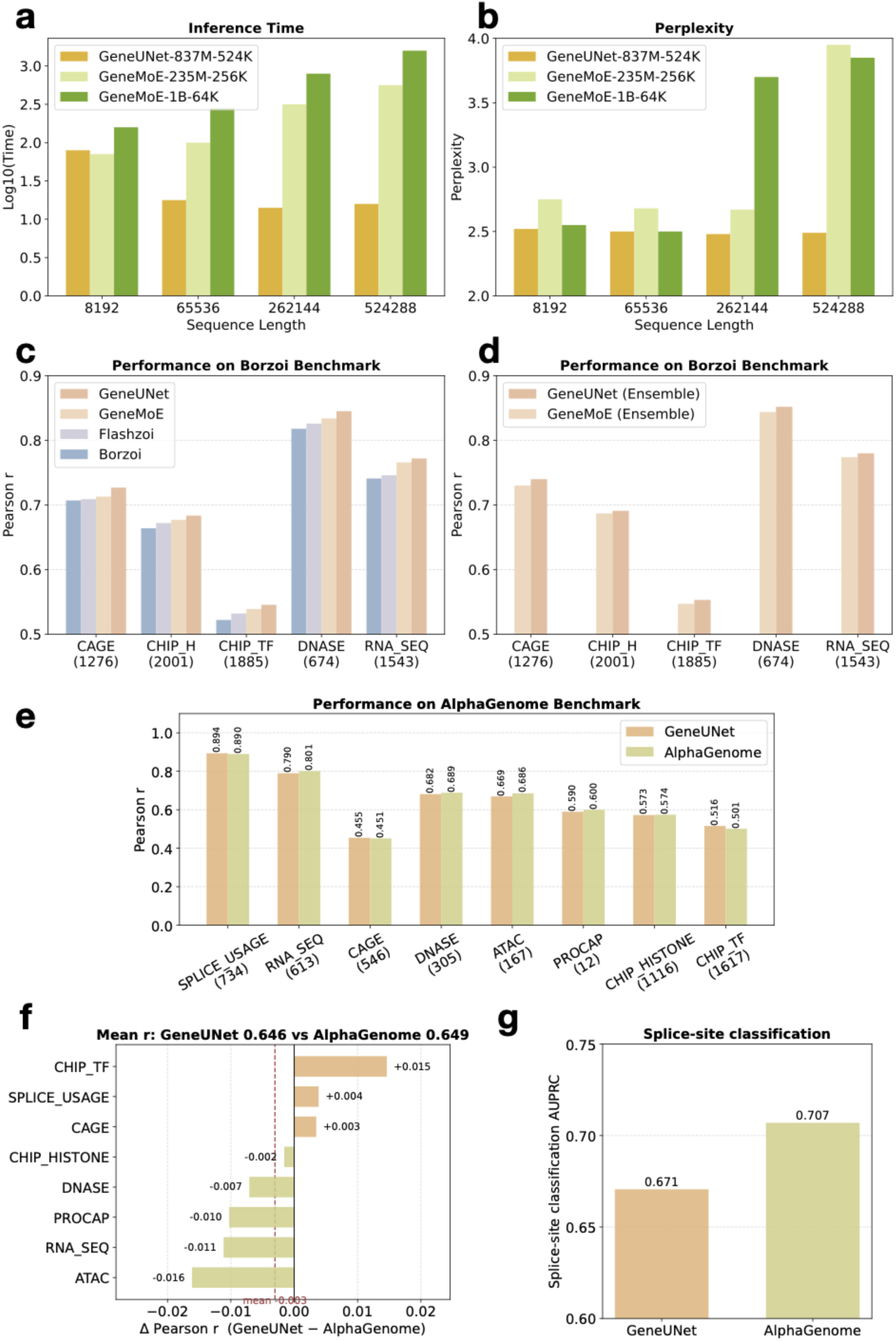
| a, Inference time across context length (8 kb to 524 kb) for GB.GeneUnet (837M parameters) and GeneMoE (235M and 1B parameters). Wall-clock time is shown on a log_10_ scale under a fixed token budget. GB.GeneUnet inference speed exceeds that of both GeneMoE variants at all tested lengths, with the gap widening monotonically. Despite carrying 3.6x more parameters than GeneMoE-235M, GB.GeneUnet is 35x faster; relative to GeneMoE-1B, GB.GeneUnet achieves approximately 100x faster inference while using only 84% of its parameters, indicating that the efficiency gain is architectural rather than parametric. **b**, Perplexity across context length for the same models. Each model reaches minimum perplexity at its pretraining context length. GB.GeneUnet maintains lower perplexity than both GeneMoE variants throughout; at 524 kb, perplexity rises for both GeneMoE models whereas GB.GeneUnet remains stable. **c**,**d**, Gene expression prediction on the Borzoi benchmark (524 kb context, 32 bp resolution). **c**, Linear-probing performance of GB.GeneUnet and GeneMoE-470M compared with the fully supervised baselines Borzoi and Flashzoi. GB.GeneUnet achieves state-of-the-art performance across all tracks, surpassing both foundation-model and supervised comparators. **d**, Ensemble performance (four replicates) of GB.GeneUnet and GeneMoE-470M. GB.GeneUnet outperforms GeneMoE-470M across all tracks, with consistent gains that generalise beyond single-model evaluation. **e**–**g**, Gene expression prediction on the AlphaGenome benchmark (1 Mb context). Fully fine-tuned GB.GeneUnet is compared with AlphaGenome across eight track categories by Pearson correlation, scored against AlphaGenome’s own released labels. Each category is evaluated in the space AlphaGenome reports it in: raw counts for splice-site usage, ATAC and DNase, and log_1p_ for the remaining five. Because log(1+*x*) is not scale invariant, our predictions are mapped into the scale of those labels before the transform (by the ratio of per-track non-zero means), while AlphaGenome’s predictions require no such mapping. Track counts per category are given in parentheses; the RNA-seq category covers the 613 ENCODE-derived tracks, the 54 GTEx-derived RNA-seq tracks being excluded because AlphaGenome did not release labels for them. All results are from fold 3 (1,576 intervals) over the central 196,608 bp window. Splice-site usage is scored over the positions that AlphaGenome’s released annotation marks as a donor or acceptor site. **e**, Per-category Pearson correlation for both models. **f**, The same comparison expressed as a per-category difference, GB.GeneUnet minus AlphaGenome. The mean over the eight categories is -0.003 (0.646 versus 0.649), and every individual difference is smaller than 0.02. **g**, Splice-site classification. This is a classification readout, scored by AUPRC and therefore reported on its own rather than averaged with the correlations: the mean over the four donor/acceptor channels, on which AlphaGenome leads (0.707 versus 0.671). Per-channel values are given in Appendix B.

On the Borzoi benchmark, we fine-tune GB.GeneUnet and GeneMoE-470M using linear probing and compare against the strongest supervised baselines, Borzoi and Flashzoi (*7*). Both pretrained models outperform these baselines, with GB.GeneUnet achieving state-of-the-art performance across all tracks (**Fig 2.c**). Under an ensemble setting with four-fold cross-validation, GB.GeneUnet maintains a consistent advantage over GeneMoE across all tracks, demonstrating robust generalizability.

On the AlphaGenome benchmark (1 Mb setting), we fine-tuned GB.GeneUnet in two phases, first training at 524 kb context and then extending to 1 Mb, and evaluate it against AlphaGenome across eight functional categories: CAGE, DNase, ATAC, RNA-seq, ChIP-TF, ChIP-Histone, PRO-cap and splice-site usage. Both models are scored on the same 1,576 held-out intervals against AlphaGenome’s own released labels, so that each is measured against an external reference rather than against our reproduction of it, and each category is evaluated in the space AlphaGenome uses for it: raw counts for splice-site usage, ATAC and DNase, and log(1+*x*) for the remaining five. The evaluation protocol is given in Appendix B and the per-category values in Appendix Table 4.

GB.GeneUnet and AlphaGenome are closely matched on this benchmark (**Fig. 2e–g, Appendix Table 4**). The unweighted mean of the eight per-category correlations is 0.646 for GB.GeneUnet against 0.649 for AlphaGenome, placing GB.GeneUnet 0.003 below AlphaGenome overall. GB.GeneUnet is ahead on TF ChIP-seq (+0.015), splice-site usage (+0.004) and CAGE (+0.003), and AlphaGenome is ahead on ATAC (0.016), RNA-seq (0.011), PRO-cap (0.010), DNase (0.007) and histone ChIP-seq (0.002). Every one of these differences is smaller than 0.02, and most are at or below the precision at which such correlations are conventionally reported, so we read the comparison as parity on this benchmark rather than as a ranking in either direction. Splice-site classification is scored by AUPRC and is reported separately, since it is a classification readout and does not belong in an average with the correlation figures. Averaged over the four donor/acceptor channels AlphaGenome reaches 0.707 against 0.671 for GB.GeneUnet, ahead on each channel individually (Appendix B).

What distinguishes GB.GeneUnet on this benchmark is the cost at which that parity is reached. Fine-tuning runs in native PyTorch on a single node of 8 H100 80 GB GPUs under standard data parallelism, without sequence parallelism, model parallelism or any other partitioning of the model across devices. Three choices make a 1 Mb-context model fit that budget. Targets are stored and shipped in a sparse (index, value) layout and scattered into the dense track-by-position matrix on the GPU, so gigabytes of zeros never cross CPU memory bandwidth. The per-interval target segments are concatenated before the loss is taken, so the full multi-track objective is evaluated in one pass over the window rather than as a separate loss per segment. Activation checkpointing is applied per component rather than to the network as a whole: each middle transformer block, each fused up-sampling and skip step, and the full-resolution final block are checkpointed individually, which is what brings a 1 Mb input within the memory of a single 8-GPU node under plain data parallelism. And the context schedule is staged, taking the majority of optimizer steps at 524 kb before extending to 1 Mb for a shorter second phase, so that the bulk of training runs where attention is cheaper and the 1 Mb phase serves to adapt the model to the longer input rather than to learn the task from scratch. Details are given in Methods and Appendix B.

## Discussion and Conclusion

GB.GeneUnet shows that multispecies pretraining and ultra-long context modelling within a Transformer-based U-Net enable improved sequence-to-function prediction. However, several limitations highlight directions for improvement.

First, the current output space remains limited. Although the model predicts regulatory tracks in multiple resolutions, it does not capture higher-order chromatin interactions such as DNA contact maps (*3*), which require modelling two-dimensional genomic structure. Extending the framework to these modalities would broaden its applicability.

Second, the use of sliding-window attention with a narrow local window constrains intermediate receptive fields, placing the burden of long-range interaction modelling on the bottleneck. This design may limit efficient propagation of distal regulatory signals. Expanding attention windows or introducing dilated attention patterns could improve multi-scale information flow.

Finally, the model is not explicitly optimized for variant effect prediction. The current objectives—masked language modelling and multi-track regression—do not directly enforce allelic sensitivity. Incorporating variant-centric benchmarks or fine-tuning on labelled datasets (for example, eQTL or saturation mutagenesis data) would be necessary to establish competitiveness in genetic variant interpretation. More broadly, extending the model toward generative sequence design represents a promising direction for leveraging long-context genomic representations.

## Methods

### GB.GeneUnet Model Architecture

GB.GeneUnet is a Transformer-based U-Net architecture with a symmetric encoder–decoder design and horizontal skip connections, operating on genomic sequences of up to 1,048,576 bp at single-base resolution (**Fig. 1.b**). The model comprises a stem, seven downsampling blocks, eight bottleneck blocks, seven upsampling blocks, and a final output block. A vocabulary of size 128 is used to encode DNA, RNA and special tokens. The stem maps input tokens to a 512-dimensional embedding, followed by a Transformer block. The final block operates at the same resolution, applying a Transformer block and a linear projection to 128 output dimensions. **Downsampling blocks** Each downsampling block consists of average pooling (stride 2), a linear projection from *d_i-_*_1_ to *d_i_*, and two Transformer blocks. A linear skip projection at dimension *d_i-_*_1_ is stored prior to pooling for use in the corresponding upsampling stage. Channel dimensions across the seven stages are [512, 768, 1024, 1280, 1536, 1792, 2048].

### Upsampling blocks

Each upsampling block applies two Transformer blocks, followed by a linear projection from *d_i_* to *d_i-_*_1_ and nearest-neighbor upsampling (scale factor 2). The corresponding skip connection from the encoder is added element-wise after upsampling. Channel dimensions follow the reverse order [2048, 1792, 1536, 1280, 1024, 768, 512].

### Transformer blocks

Transformer blocks in the encoder and decoder employ sliding-window flash attention (SWA) (*8–10*) with a one-sided window of 4 tokens (maximum context of 8 neighbouring positions), grouped-query attention (GQA) (*11*) with *n*_kv_ = *n*_head_*/*2, and rotary positional embeddings (RoPE) with base frequency 100,000. The number of attention heads *n_head_* across downsampling blocks is [8, 8, 8, 8, 8, 16, 16] (reversed in upsampling). The feed-forward sub-layer uses SwiGLU with expansion ratio 4, projecting from *d* to 4*d*, splitting into two halves with SiLU gating, and projecting back to *d*.

### Bottleneck blocks

Bottleneck blocks follow the same design as the standard Transformer blocks, with three modifications. First, attention is computed globally over all tokens, without window restriction. Second, GQA uses 16 query heads and 8 key-value heads. Third, the feed-forward sub-layer adopts a two-layer GELU network with an expansion ratio of 2 instead of SwiGLU.

### Pretraining

#### Pretraining Data

Following Evo2 (*6*), we pretrain GB.GeneUnet on the OpenGenome2 database, which comprises approximately 8.8 trillion base pairs of curated DNA spanning all domains of life. The dataset includes 357 billion nucleotides from prokaryotic genomes, 6.98 trillion nucleotides from eukaryotic genomes, 854 billion nucleotides of non-redundant metagenomic sequences, 2.82 billion nucleotides from organelle genomes, and 602 billion nucleotides from functionally annotated eukaryotic genomic regions, including mRNA and non-coding RNA (ncRNA) transcripts, non-coding RNAs, and eukaryotic promoter regions.

OpenGenome2 is processed differently during the pretraining and midtraining phases. During pretraining, data augmentation and filtering are applied to emphasize conserved and information-dense regions surrounding genes. In contrast, during midtraining, the effective sequence length is substantially increased for prokaryotic and eukaryotic genomes, allowing the model to observe long-range genomic dependencies within a single training sequence.

Specifically, during pretraining, 50% of sequences are randomly reverse complemented to improve robustness to strand orientation. mRNA and ncRNA transcript sequences are augmented by extension with upstream promoter regions and exon-centered flanking sequences to capture splice-site context, and by stitching transcript components into composite sequences. In addition, exon-centered genomic windows are extracted from eukaryotic genomes to enrich the training data with coding and cis-regulatory context.

During midtraining, transcript-based and augmented datasets remain unchanged, while genome-scale sequences from the same accession are concatenated to yield effective sequence lengths on the order of millions of base pairs, enabling the model to learn long-range dependencies across extended genomic regions.

#### Pretraining Setting

The model is pretrained on genomic sequences from OpenGenome2 using a masked language modelling (MLM) objective with a masking ratio of 15%, optimized via cross-entropy loss (**Fig. 1.a**). Pretraining is conducted in two consecutive stages with identical training and optimization settings. In the first stage, the model is trained on 3 trillion tokens from the OpenGenome2 pretraining phase. In the second stage, training is resumed from the first-stage checkpoint and continued for an additional 3 trillion tokens using data from the OpenGenome2 midtraining phase.

Training is performed on 8 NVIDIA H100-80G GPUs using data parallelism. We used the AdamW optimizer with a learning rate of 1 × 10*^-^*^4^ and weight decay of 0.1, a 1,000-step linear warmup followed by linear decay over the final 10% of training. Mixed-precision training (bfloat16) and gradient checkpointing are enabled. Each sequence is randomly reverse-complemented with probability 0.5, with strand-paired tracks permuted accordingly as data augmentation.

### Finetuning for Borzoi Benchmark

GB.GeneUnet and GeneMoE-medium are benchmarked against Borzoi (*2*) and its efficient variant Flashzoi (*7*) on the Borzoi benchmark. Following the original data partitioning scheme (*2*), both human and mouse datasets are divided into eight non-overlapping folds; folds 0–2 and 5–7 serve as the training set, fold 4 as validation, and fold 3 as the held-out test set. During training, input sequences and their corresponding coverage tracks are randomly reverse-complemented, and sequences are randomly shifted by 0–3 nucleotides without altering the labels, following standard data augmentation practices.

Frozen GB.GeneUnet and GeneMoE encoders are coupled to a Flashzoi adapter to predict multiple tracks expression from 524-kb input sequences. The convolutional DNA stem of Flashzoi is replaced by a linear projection that maps encoder hidden states to 512 channels, matching the expected input dimensionality of the Flashzoi trunk. To accommodate long sequences efficiently, sliding window attention with a local context of 8,192 tokens is applied within the frozen encoder. Adapter training uses bfloat16 mixed precision on 32 H100 GPUs (global batch size = 32), with a linear warm-up over 2,000 steps to a peak learning rate of 2 x 10^-5^, weight decay of 1 x 10^-8^, and gradient clipping at 0.15. Early stopping is applied when validation Pearson correlation coefficient (PCC) shows no improvement for 30 consecutive evaluations.

For ensemble predictions, four Flashzoi adapters are independently initialized from distinct publicly available checkpoints (johahi/flashzoi-replicate-{0,1,2,3}) with randomized sample orderings, and their outputs are averaged.

### Finetuning for AlphaGenome Benchmark

We combined the pretrained GB.GeneUnet with two lightweight species-specific adapters for human and mouse, and fine-tuned the full model to predict multi-track epigenomic and transcriptomic signals at 1 bp resolution (ATAC-seq, CAGE-seq, DNase-seq, PRO-cap, RNA-seq, splice-site usage) and 128 bp resolution (ChIP-histone and ChIP-TF) in two phases on 8 H100 80GB GPUs. Splice supervision changed source during the project. We first trained on splice targets of our own construction: four donor/acceptor channels derived from GENCODE gene annotations (*12*) together with GTEx v8 splice-junction data accessed via recount3 (*13*), and splice-site-usage tracks derived from ENCODE, under the sources and terms described below. Having identified defects in that preparation, we then ran a short continued-training phase in which both the splice-site and the splice-site-usage targets were replaced by AlphaGenome’s released labels (*3*), supplied at data-loading time so that no stored target is modified; the model reported here is the result of that phase. Both splice readouts are therefore evaluated against AlphaGenome’s released labels: splice-site usage by Pearson correlation, alongside the other seven categories, and splice-site classification by AUPRC, which is reported separately and is not averaged into the correlation figures. The fine-tuning targets were derived from the original public data sources, processed following the AlphaGenome data-processing methodology (*3*): ENCODE (ATAC-seq, DNase-seq, PRO-cap, RNA-seq, ChIP-seq, and splice tracks) under the ENCODE Data Use Policy (*14*); FANTOM5 (CAGE tracks) under the Creative Commons Attribution (CC-BY) license (*15*); GTEx v8 (a subset of RNA-seq and splice-site tracks), accessed via recount3 (*13*) under the GTEx General Research Use terms (*16*); and GENCODE gene annotations under the EMBL-EBI terms of use (*12*). A subset of splice-site-usage tracks additionally incorporated the corresponding ENCODE-derived records from the released Al-phaGenome training data (*3*), used under the ENCODE Data Use Policy (*14*). The four splice-site classification channels (donor and acceptor on each strand) were derived from GENCODE gene annotations (*12*) together with GTEx v8 splice-junction data accessed via recount3 (*13*), under the GTEx General Research Use terms (*16*); these are the same sources from which AlphaGenome’s corresponding splice-site data are processed. The splice-site-usage tracks are ENCODE-derived and do not draw on GTEx. Consistent with GTEx General Research Use terms, GTEx-derived ground-truth data are not included in any externally released or shared datasets. Training used human and mouse folds 0, 1, 2, 5, 6, and 7, validation used human fold 4, and testing used the held-out human fold 3, following the Borzoi fold-split convention (*2*)^1^ (**Fig. 1.a**). AlphaGenome baseline scores in Table 4 were obtained from the AlphaGenome API (*3*) using its fold-1 model rather than an all-folds model, so that fold 3 lies outside its training data, and were evaluated on the fold-3 test intervals with the same pipeline used for GB.GeneUnet. All scores reported in Table 4 are computed against AlphaGenome’s own released labels for those intervals, so that both models are measured against an external reference rather than against our reproduction of the targets; because log(1+*x*) is not scale invariant, our predictions are mapped into the scale of those labels before the transform, while AlphaGenome’s require no such mapping. Reverse-complement augmentation is applied during training, with a coordinate shift on stranded DNase tracks to preserve strand-pair alignment. The 128 bp ChIP heads are placed on the central (low-resolution) bottleneck of the U-Net, and a per-track learnable scale is applied to all head outputs to absorb track-specific magnitude differences. The full backbone is unfrozen for fine-tuning.

**Phase 1.** Training is conducted at a context length of 524,288 bp for approximately 25,000 optimizer steps. The Muon optimizer is applied to the backbone parameters with a peak learning rate of 1.2 × 10*^-^*^3^. AdamW is used for the rest of the parameters: the prediction heads with a peak learning rate of 1 × 10*^-^*^3^ and the adapter modules with a peak learning rate of 1 × 10*^-^*^4^. The schedule is a linear decay over the full training horizon with no warmup. Weight decay is 10*^-^*^8^ and gradient norms are clipped at 0.1. Per-track loss weights are set to 1.0 for all track types except PROCAP, which is upweighted to 5.0 to compensate for its underrepresentation. Targets are stored on disk as 131,072 bp segments; at 524 kb context the 4 central segments of an interval are loaded in sparse form and concatenated on the GPU into a single contiguous target window, so the objective is evaluated once over the whole window rather than once per segment. Each rank processes one such sample (micro-batch of 1), and gradients are accumulated across ranks to reach the global batch of 32. The best checkpoint by validation Pearson on human tracks is selected for Phase 2.

**Phase 2 (context extension to 1 Mb).** The context length is extended from 524 kb to 1,048,576 bp for a further 20,000 optimizer steps, initialized from the Phase 1 checkpoint with the optimizer state reset. To accommodate the 2× context extension, NTK-aware RoPE scaling is applied by overriding the middle-layer RoPE base from 100,000 to 200,000. The same segment assembly is used, with the 6 central segments loaded and concatenated per sample. The training loss is computed on the central 720,896 bp of the model output (`model_crop_size`=163,840 bp on each side of the 1 Mb input); at evaluation time, predictions are further cropped to the central 196,608 bp so that the eval window is identical to Phase 1 and the AlphaGenome benchmark. A ±10 bp random shift is applied to input coordinates as further augmentation. Learning rates inherit the Phase 1 peak values (1.2 × 10*^-^*^3^ Muon for the backbone, 1 × 10*^-^*^3^ AdamW for the heads, 1 × 10*^-^*^4^ AdamW for the adapter), decayed linearly to zero over the 20,000 steps with no warmup. Weight decay, gradient clipping, per-track loss weights, RC augmentation, ChIP head placement, and per-track scale are all inherited unchanged from Phase 1.

During training, sequences are randomly reverse-complemented with probability 0.5, with strand-paired tracks permuted accordingly. For DNase-seq tracks, an additional 1 bp shift correction is applied after reverse complementation to account for the asymmetric cut-site convention. No data augmentation is applied during validation or testing. The 5x weighting of PROCAP from phase 1 is preserved.

## Data and Code Availability

GB.GeneUnet prediction scores on the AlphaGenome benchmark test set, the corresponding AlphaGenome-released ground-truth labels against which they are scored, and downstream evaluation scripts are publicly available at https://drive.google.com/drive/folders/1rhXF2haFnz76JchEgQiqJagWrY7N1v4K?usp=sharing.

An inference API for interactive visualization and evaluation of GB.GeneUnet predic-tions is accessible at https://gorgeous-cinnamon-spinner.ngrok-free.dev. As GB.GeneUnet is currently under commercial development by GenBio AI, model weights are not publicly available at this stage.

## Footnotes

1 AlphaGenome’s fold-1 release assigns the two held-out Borzoi folds to the opposite roles: fold 3 is its validation fold and fold 4 its test fold, whereas the Borzoi convention we follow takes fold 3 as the test fold and fold 4 as validation. Both models are therefore trained on the same six folds (0, 1, 2, 5, 6 and 7), and fold 3 is outside the training data of both. AlphaGenome confirmed this swap, and confirmed that no model selection was performed on either held-out fold, in response to a question on their community forum: “It is possible that the validation and test set were swapped between Borzoi and AlphaGenome (fold 3 vs fold4)” and “The fold1 model was trained without any consideration of the metrics in either validation or test set (we also haven’t used any early stopping)” (*17*). Fold 3 is thus a valid held-out test set for both systems: neither trained on it, and neither used it for checkpoint selection.

