## Supplementary for "Pretraining Enhances Megabase-Scale Gene Expression Prediction with GeneUnet"

### A GeneMoE Model

GeneMoE is a mixture-of-experts (MoE)-based DNA foundation model pretrained on the same dataset as GB.GeneUnet using masked language modelling (MLM) with cross-entropy loss. To promote computational balance and training stability across experts, load-balancing loss and router z-loss were incorporated during pretraining. Three model variants were developed spanning a range of scales: 235 million with 256K context length, 470 million with 64K context length, and 1 billion parameters with 64K context length; architectural details—including the number of layers, hidden size, and feed-forward network (FFN) dimensions—are provided in Table 2, and stage-wise training hyperparameters are summarized in Table 3.(**Fig 1**).

#### A.1 MoE Architecture

GeneMoE employs a MoE-based transformer encoder architecture, building upon the encoder of vanilla Transformer. The following improvements have been implemented: (1) Pre-LayerNorm is adopted instead of Post-LayerNorm; (2) FlashAttention-3 is utilized to enhance computational efficiency; (3) Rotary Positional Encoding (RoPE) is incorporated for position representation; (4) RMSNorm with zero-centered gamma is used in place of LayerNorm; (5) The standard Feed-Forward Network (FFN) is replaced with a Mixture of Experts (MoE); (6) The SwiGLU activation function is applied.

Within the MoE component, the architecture consists of 8 routed experts and 1 shared expert. Each token is routed to 2 routed experts. Each routed expert incorporates a learnable bias, which is added to the router's output weights to facilitate top-$K$ expert selection. Let $\mathbf{u}_t$ be the FFN input of the $t$-th token; we compute the FFN output $\mathbf{h}'_t$ as follows:

$$\mathbf{h}'_t = \mathbf{u}_t + \sum_{i=1}^{N_s} \mathrm{FFN}_i^{s}(\mathbf{u}_t) + \sum_{i=1}^{N_r} g_{i,t}\, \mathrm{FFN}_i^{r}(\mathbf{u}_t) \tag{1}$$

$$g_{i,t} = \begin{cases} s_{i,t}, & s_{i,t} \in \mathrm{TopK}(\{s_{j,t}\}_{1 \le j \le N_r}, K_r) \\ 0, & \text{otherwise} \end{cases} \tag{2}$$

$$s_{i,t} = \mathrm{Sigmoid}\left(\mathbf{u}_t^{\top} \mathbf{e}_i + b_i\right) \tag{3}$$

where $N_s$ and $N_r$ denote the numbers of shared experts and routed experts, respectively; $\mathrm{FFN}_i^{s}(\cdot)$ and $\mathrm{FFN}_i^{r}(\cdot)$ denote the $i$-th shared expert and the $i$-th routed expert, respectively; $K_r$ denotes the number of activated routed experts; $g_{i,t}$ is the gate value for the $i$-th expert; $s_{i,t}$ is the token-to-expert affinity; $\mathbf{e}_i$ is the centroid of the $i$-th routed expert in this layer; and $\mathrm{TopK}(\cdot, K)$ denotes the set comprising $K$ highest scores among the affinity scores calculated for the $t$-th token and all routed experts.

#### A.2 Pretraining Loss

**Cross-Entropy Loss** We use masked language modeling to train the model: 15% of input tokens are selected; 80% among them are replaced with `[MASK]`, 10% are replaced with random tokens, and the remaining 10% are kept unchanged. The cross-entropy loss is:

$$\mathcal{L}_{\mathrm{ce}} = -\sum_{i \in \mathcal{M}} y_i \log(p_i) \tag{4}$$

where $\mathcal{M}$ is the set of masked tokens, $y_i$ is the label, and $p_i$ is the model probability on the true token.

**Load Balancing Loss** We use an expert-level balance loss as in DeepSeek-V2 to mitigate the risk of routing collapse:

$$\mathcal{L}_{\mathrm{ExpBal}} = \sum_{i=1}^{N_r} f_i \cdot P_i \tag{5}$$

$$f_i = \frac{N_r \cdot K_r}{T} \sum_{t=1}^{T} \mathbf{1}(\text{Token } t \text{ selects Expert } i) \tag{6}$$

$$P_i = \frac{1}{T} \sum_{t=1}^{T} s_{i,t} \tag{7}$$

where $\mathbf{1}(\cdot)$ denotes the indicator function and $T$ denotes the number of tokens in a sequence.

**Z-Loss** We also use router z-loss to improve training stability (following ST-MoE), defined as:

$$\mathcal{L}_z = \frac{1}{T} \sum_{t=1}^{T} \left( \log \sum_{j=1}^{N_r} e^{x_j^{(t)}} \right)^2 \tag{8}$$

$$x_j^{(t)} = \mathbf{u}_t^\top \mathbf{e}_j \tag{9}$$

where $T$ is the number of tokens, $N_r$ is the number of experts, and $x_j^{(t)} \in \mathbb{R}^{T \times N_r}$ are the logits going into the router.

The final loss is:

$$\mathcal{L} = \mathcal{L}_{\text{ce}} + \alpha_1 \, \mathcal{L}_{\text{ExpBal}} + \alpha_2 \, \mathcal{L}_z \tag{10}$$

where $\alpha_1$ and $\alpha_2$ are hyperparameters called the expert-level balance loss coefficient and the MoE z-loss coefficient, respectively.

#### A.3 Token-Dropping

We also introduced a device-level token-dropping strategy during training, following DeepSeek-V2. The computation budget $C$ for each device is pre-calculated as:

$$C = \frac{T \cdot K_r}{N_r} \cdot C_r \tag{11}$$

The capacity factor $C_r$ for each device is set to $1.125$. If the number of routed tokens for a certain device exceeds $C$, random tokens are dropped so that the maximum number of processed tokens per device is $C$.

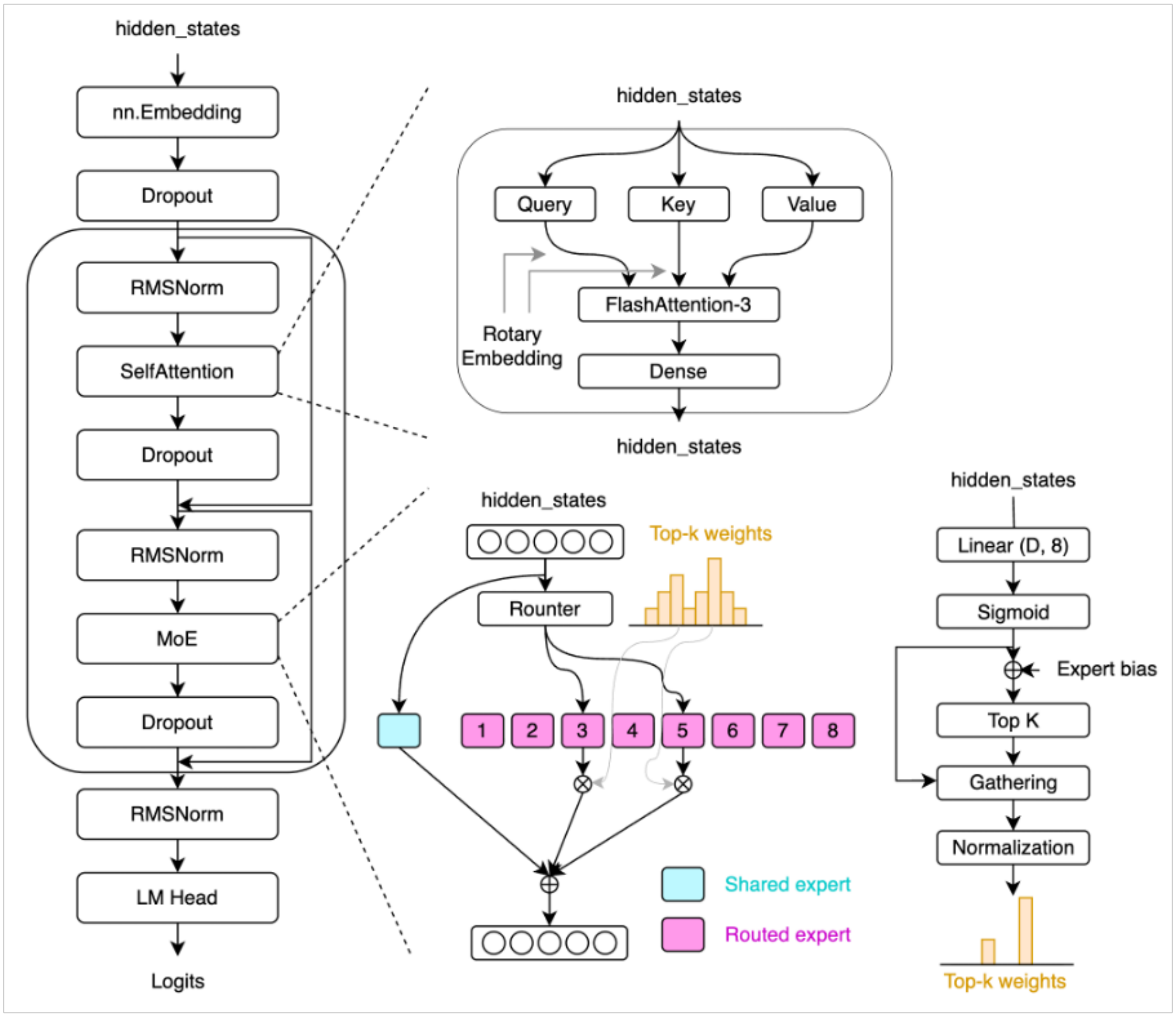


Figure 1 | GeneMoE model architecture

### B AlphaGenome benchmark: protocol and per-category results

**Interval set.** All $1{,}576$ human intervals of Borzoi fold 3 are used. Fold 3 is outside the training data of both GB.GeneUnet and AlphaGenome (see the fold-assignment footnote in Methods). Predictions and labels are cropped to the central $196{,}608$ bp of each interval. AlphaGenome's released windows are $1{,}052{,}672$ bp wide and ours are $1{,}048{,}576$ bp; the two are paired by shared genomic centre, not by filename.

**Labels.** Both models are scored against AlphaGenome's own released fold-3 labels, so that neither is measured against our reproduction of the targets. AlphaGenome predictions were obtained from the AlphaGenome API using its fold-1 model rather than an all-folds model, so that fold 3 lies outside its training data.

**Metric.** Pearson correlation is computed independently for each track over the concatenation of all intervals, then averaged arithmetically over the tracks of a category (`nanmean`: tracks whose label or prediction is constant across the whole test set are undefined and excluded). This is the aggregation used in the original AlphaGenome evaluation. Because $\log(1{+}x)$ is not scale invariant, our predictions are mapped into the scale of the released labels before the transform; AlphaGenome's predictions require no such mapping.

**Spaces.** Each category is evaluated in the space AlphaGenome uses for it: raw counts for splice-site usage, ATAC and DNase, and $\log(1{+}x)$ for CAGE, RNA-seq, PRO-cap, histone ChIP-seq and TF ChIP-seq.

**Splice-site usage.** Usage is defined only at splice sites, so the correlation is taken over the positions that AlphaGenome's released splice-site annotation marks as a donor or an acceptor (i.e.

any non-background channel), pooled over all intervals. This is the evaluable-position definition used in the original evaluation. AlphaGenome places its splice outputs on the flanking exonic bases (donor at $p - 1$, acceptor at $p + 1$, relative to the intron-terminal base $p$ used by STAR), and GB.GeneUnet's splice head is trained in the STAR frame, so GB.GeneUnet's splice-usage predictions are shifted by one base with a class-conditional offset before scoring. AlphaGenome is scored against its own labels with no shift applied.

**RNA-seq track count.** AlphaGenome's released 1 bp axis carries 613 RNA-seq tracks plus 54 padding columns standing in for its unreleased GTEx RNA-seq tracks. RNA-seq is therefore scored over the 613 released ENCODE tracks.

Averaged over the eight categories the two fold-3 rows differ by 0.003 in Pearson correlation. GB.GeneUnet leads on TF ChIP-seq (+0.0147), splice-site usage (+0.0038) and CAGE (+0.0034), and is within 0.002 of AlphaGenome on histone ChIP-seq (−0.0016). AlphaGenome leads on ATAC (−0.0161), RNA-seq (−0.0112), PRO-cap (−0.0102) and DNase (−0.0071). Expressed as a relative difference per category and averaged, GB.GeneUnet is 0.33 % below AlphaGenome; taking the ratio of the two means instead gives 0.47 %.

**Splice-site classification.** The four donor/acceptor channels are a classification readout and are scored by AUPRC rather than by correlation, computed per channel over all positions of all 1,576 intervals against AlphaGenome's released splice-site annotation, then averaged over the four channels. AUPRC is not averaged into the Pearson figures of Table 4. The histograms backing these values use 65,536 score bins, which is exact here because the stored predictions are `float16`.

AlphaGenome is ahead on every channel, and by 0.036 on the mean.

Table 1 | Splice-site classification AUPRC on the same 1,576 fold-3 intervals, per donor/acceptor channel and averaged over the four. Both models are scored against AlphaGenome's released splice-site annotation. Bold marks the better of the two.

| | donor+ | acceptor+ | donor− | acceptor− | Mean |
|---|---|---|---|---|---|
| GB.GeneUnet | 0.6795 | 0.6552 | 0.6611 | 0.6868 | 0.6707 |
| AlphaGenome | **0.7165** | **0.6908** | **0.6947** | **0.7259** | **0.7070** |

Table 2 | Architectural details of GeneMoE model variants.

| | **GeneMoE-235M** | **GeneMoE-470M** | **GeneMoE-1B** |
|---|---|---|---|
| Layers | 8 | 16 | 16 |
| Hidden size | 1024 | 1024 | 1536 |
| Number of routed experts | 8 | 8 | 8 |
| Number of shared experts | 1 | 1 | 1 |
| Number of selected experts | 2 | 2 | 2 |
| FFN hidden size of routed experts | 512 | 512 | 768 |
| FFN hidden size of shared experts | 4096 | 4096 | 6144 |
| Number of heads | 8 | 8 | 8 |
| Activated / Total parameters (M) | 159 / 235 | 319 / 470 | 717 / 1056 |

#### Additional Technical Details on GB.GeneUnet training for the Alphagenome Task

**Training budget and hardware** Both models are trained on a single node with 8× NVIDIA H100 80 GB GPUs, using PyTorch with `bfloat16` mixed precision and standard data-parallel training (DDP). The activation-checkpointing and label-handling strategies described below and in Fig. 2) are what allow us to fit a 1 Mb-input model into a single 8-GPU node with DDP, rather than relying on sequence parallelism or other parallelism technique. The 524 kb phase is trained for 143 wall-clock hours ($\approx$ 6 days) with a global batch size of 32 (micro-batch of 1 per rank, gradient accumulation across ranks). GB.GeneUnet is obtained by fine-tuning from the Phase 1 checkpoint with the optimizer state reset, and trains for 22 further hours. Both figures cover the two context phases only; they exclude the short continued-training phase on AlphaGenome's

Table 3 | Training hyperparameters for each stage of GeneMoE pretraining.

| | **Models** | **Stage 1** | **Stage 2** | **Stage 3** | **Stage 4** |
|---|---|---|---|---|---|
| Dataset | All | OG2-pre | OG2-pre | OG2-mid | OG2-mid |
| Sample length | All | 2,048 | 8,192 | 64,000 | 256,000 |
| Micro / Global batch size | GeneMoE-235M | 32 / 4096 | 8 / 768 | 1 / 96 | 1 / 32 |
| | GeneMoE-470M | 32 / 4096 | 8 / 768 | 1 / 96 | N/A |
| | GeneMoE-1B | 32 / 4096 | 4 / 768 | 1 / 96 | N/A |
| Learning rate schedule (WSD) | GeneMoE-235M | 2e-6 $\sim$ 1e-4 | 2e-6 $\sim$ 1e-4 | 0 $\sim$ 5e-5 | 0 $\sim$ 5e-5 |
| | GeneMoE-470M | 2e-6 $\sim$ 1e-4 | 2e-6 $\sim$ 1e-4 | 0 $\sim$ 5e-5 | N/A |
| | GeneMoE-1B | 2e-6 $\sim$ 1e-4 | 2e-6 $\sim$ 1e-4 | 0 $\sim$ 5e-5 | N/A |
| Training tokens | All | 750B | 250B | 250B | 100B |
| Number of nodes | GeneMoE-235M | 4 | 4 | 4 | 4 |
| | GeneMoE-470M | 4 | 4 | 4 | N/A |
| | GeneMoE-1B | 4 | 4 | 6 | N/A |
| TP/PP/CP/EP | GeneMoE-235M | 1/1/1/8 | 1/1/1/8 | 1/1/1/8 | 1/1/1/8 |
| | GeneMoE-470M | 1/1/1/8 | 1/1/1/8 | 1/1/1/8 | N/A |
| | GeneMoE-1B | 1/1/1/8 | 1/1/1/8 | 1/1/1/8 | N/A |

released splice labels described in Methods, from which the reported model is taken, so the total cost of the reported model is somewhat higher.

**Muon optimizer and parameter partitioning** Following recent work on matrix-adapted optimizers (*?*), we use a hybrid scheme that combines Muon for matrix-shaped parameters with AdamW for the rest, implemented using the `MuonWithAuxAdam` class from the original Muon github repository from Keller Jordan. Parameter groups are selected by two criteria computed from the parameter's fully-qualified name and tensor rank:

- **Backbone matrix parameters** (with `ndim` $\geq$ `2`: Muon group, learning rate 12x higher than AdamW, momentum 0.95, weight decay $10^{-8}$. These are the weights of every linear / QKV / MLP projection inside the down-, middle-, and up-path transformer blocks.

- **Backbone 1D parameters** (`backbone.*` with `ndim = 1`): AdamW, $\beta_1$=0.9, $\beta_2$=0.999, no weight decay. These are the RMSNorm scales and any remaining biases.

- **Head matrix parameters** (`adapter.*` or `splice_head.*` with `ndim` $\geq$ `2`): AdamW, learning rate $7.29 \times 10^{-5}$, weight decay $10^{-8}$.

- **Head 1D parameters**: AdamW, learning rate $7.29 \times 10^{-5}$, no weight decay.

This partition is done automatically from `self.named_parameters()`: every parameter is routed by backbone-vs-head namespace and by tensor rank, so no per-layer configuration is required when the architecture changes. The Muon group covers the bulk of the model parameters ($\approx$ 95% of the parameter count in our 837M configuration), where we find Muon's matrix-orthogonalized updates to meaningfully outperform AdamW at fixed wall-clock; the remaining 1D buffers stay on AdamW because they are scalar- per-output and do not benefit from the Newton–Schulz orthogonalisation.

**Context extension and rotary embeddings** To double the input context from 524 288 bp to 1 048 576 bp without retraining the attention positional representations from scratch, we apply NTK-aware rotary-embedding scaling in the middle transformer blocks: the RoPE base frequency is increased from $1.0 \times 10^5$ to $2.0 \times 10^5$, a factor of 2 matched to the 2$\times$ longer context. This preserves the effective angular frequencies at integer positions while extending the unambiguous position range, so the model's position-dependent attention behaviour is a smooth extrapolation of what it learned at 524 288 bp, and fine-tuning converges in $\approx 10^4$ optimizer steps rather than requiring full retraining.

**Output cropping and matched label windows** GB.GeneUnet internally crops its 1 bp and 128 bp prediction heads to a central sub-window of the input. At 1 Mb input we default to predicting the central 196 608 bp (a symmetrical 425 984 bp trim on each side), makes GB.GeneUnet directly comparable to the 524 kb phase on the same target region. Restricting supervision to the central window is motivated by two observations: (i) predictions near the edges of the input see attenuated attention context (the full $\pm 500$ K surround is only available at the centre), so edge predictions are systematically noisier; and (ii) computing loss and metrics on the centre only reduces peak activation memory and wall-clock per step, and in our experiments did not degrade validation quality relative to supervising a wider window — an important enabler for fitting 1 Mb context into a single 8-GPU node.

**Activation checkpointing** Both models use activation checkpointing: intermediate activations are freed on the forward pass and rematerialised on the backward pass, exchanging compute for memory (see Fig. 2 for a side-by-side comparison of the two checkpointing configurations).

**Phase 1 (524 kb).** Down-sampling and up-sampling layers are each individually checkpointed, and the entire middle transformer stack is wrapped in a single checkpoint. This is sufficient at 524 288 bp input on 80 GB H100s.

**Phase 2 (1 Mb).** Doubling the context to 1 048 576 bp doubles the activations per middle block, and under the Phase 1 strategy the 1 Mb model would exceed the 80 GB H100 capacity. We therefore apply four additional memory optimizations:

1. **Per-block middle checkpointing.** Each transformer block in the middle stack is wrapped in its own checkpoint, rather than the whole stack being a single checkpoint. Peak rematerialisation memory during backward reduces from "all blocks at once" to "one block at a time".

2. **Deferred horizontal skip.** The U-Net horizontal skip convolutions are not evaluated on

the down path; instead, lightweight references to the down-layer inputs are retained, and the horizontal features are recomputed inside a combined checkpoint on the up path. This avoids keeping multiple full-resolution skip tensors alive across the forward pass.

3. **Combined up + skip checkpoint.** Each up-sampling layer is fused with its matching horizontal convolution inside a single checkpoint, so the horizontal activation is never materialised outside the checkpoint boundary ($\approx 3\,\text{GB}$ saved per up step).

4. **Final-block checkpoint.** At 1 Mb resolution the final transformer block (attention + feed-forward at full length) uses $\approx 25\,\text{GB}$ of intermediates without checkpointing; we wrap it in a checkpoint as well.

On top of the backbone, both runs additionally checkpoint the per-track-type adapter heads during training, trading a constant memory reduction for $2\times$ adapter recompute. FlashAttention-3 is used in every middle self-attention layer.

Together, these checkpointing techniques, combined with the matched output window described above, allow us to fit GB.GeneUnet on a single 8-GPU node under standard PyTorch DDP; we do not require sequence parallelism or any multi-node partitioning of the attention matrices.

**Sparse target handling** Most 1 bp labels (CAGE, PROCAP, splice sites, ATAC, DNase peaks) are exactly zero at most positions, so we store them in a sparse (nonzero-index, nonzero-value) layout rather than dense. Dense labels are never materialised on the CPU: the dataloader ships the sparse tensors to the GPU, where a single scatter operation reconstructs the dense [tracks $\times$ positions] matrix. This avoids moving gigabytes of zeros through CPU memory bandwidth every step and lets the label reconstruction overlap with the forward pass on the previous batch.

**Reverse-complement and shift augmentation** Training applies reverse-complement (RC) augmentation with probability 0.5 and a random 1 bp shift $s \in \{-1, 0, +1\}$. Two subtleties deserve attention.

**DNase shift.** DNase-seq signal is recorded at the 5' end of each cleaved fragment. Reversing the DNA moves the effective 5' position by 1 bp, and we compensate by shifting the DNase label by $+1$ along the sequence axis after the RC flip. Without this correction, RC-augmented training biases the model's DNase predictions by $\pm 1$ bp.

**Strand-specific track permutation.** Stranded readouts have to be re-mapped after RC: the RNA-seq$+$ channel on the original reference becomes the RNA-seq signal from the opposite strand, i.e. the original RNA-seq$-$. We therefore swap every stranded-track pair when RC is applied (CAGE$\pm$, RNA-seq$\pm$, PROCAP$\pm$, splice-site usage $\pm$, and the splice-site class channels), with strand-paired channels swapped accordingly.

**Validation and test split** Following the standard Borzoi convention, fold 3 is held out as the test fold and chromosome fold 4 as the validation fold; training iterates over folds $\{0, 1, 2, 5, 6, 7\}$. Running validation during training is computed on a fixed 10% ($\approx 158$) fold-4 intervals. Final test metrics are reported on the full 1 576 fold-3 intervals, evaluated on the central 196 608 bp of the 1 Mb input for both training phases, at 1 bp resolution for regression / SSU / SS classification and at 128 bp resolution for CHIP-seq signals.

**Effect of pretraining on the AlphaGenome task.** To isolate the contribution of multi-species pretraining, we compare two 524 kb fine-tuning runs that share an architecture, dataset and training recipe but differ only in initialization: one initialized from the OpenGenome2 pretrained checkpoint ("Pretrained"), and one with all weights reinitialized from scratch ("From scratch"). Both are fine-tuned on the AlphaGenome task under identical DDP, optimizer and augmentation settings. Figure 3 reports the per-track validation Pearson correlation on human fold 4 over the

Table 4 | GB.GeneUnet versus AlphaGenome on the AlphaGenome benchmark, human Borzoi fold 3 ($1{,}576$ intervals, central $196{,}608$ bp). All values are Pearson correlation, computed per track and averaged over the tracks of a category. Following the original AlphaGenome evaluation, CAGE, RNA-seq, PRO-cap, ChIP-hist. and ChIP-TF counts are $\log(1{+}x)$-transformed, while splice-site usage, ATAC and DNase are evaluated on raw counts. Bold marks the better of the two fold-3 rows; where both are bold they agree to two decimal places. For reference we also quote AlphaGenome's published results, read from the violin plot in main Figure 2c of (?); these are computed on AlphaGenome's own held-out test split rather than on fold 3, and are excluded from the bolding. See Appendix B for the evaluation protocol.

| | Splice usage | CAGE | RNA-seq | ATAC | DNase | PRO-cap | ChIP-hist. | ChIP-TF |
|---|---|---|---|---|---|---|---|---|
| GB.GeneUnet (fold 3) | **0.89** | **0.45** | 0.79 | 0.67 | 0.68 | 0.59 | **0.57** | **0.52** |
| AlphaGenome (fold 3) | **0.89** | **0.45** | **0.80** | **0.69** | **0.69** | **0.60** | **0.57** | 0.50 |
| AlphaGenome (published) | 0.86 | 0.46 | 0.81 | 0.70 | 0.67 | 0.60 | 0.59 | 0.52 |

first $80{,}000$ optimizer steps. The pretrained model reaches a higher plateau on every evaluated track, and the gap is established within the first few thousand steps rather than appearing only at convergence; aggregated across tracks, the pretrained model is ahead by $\approx 0.020$ on the overall Pearson at $80{,}000$ steps (Table 5). We note two caveats specific to this experiment. First, these runs used an earlier version of the AlphaGenome dataset (pre-patched splice labels, older PROCAP weighting and augmentation settings) and a training recipe that differs from the main-text main-text runs; absolute scores are therefore not directly comparable to the fold-3 test numbers reported elsewhere in the paper. Second, the Pearson correlation shown here is computed in the model's re-scaled (raw) output space, not in $\log(1{+}p)$ space. Despite these differences, the two runs were launched back-to-back with matched settings, so the pretrained-vs.-scratch comparison *within* this figure is self-consistent and the gap is attributable to initialization.

**Signal scale.** Every correlation reported in this work is computed against AlphaGenome's released labels, in the scale of those labels. Our predictions are produced in the scale of our own reproduced targets and are therefore mapped into AlphaGenome's scale before the $\log(1{+}x)$ transform, by the ratio of per-track non-zero means; AlphaGenome's predictions require no such

Table 5 | **Pretraining vs. random initialization at $\approx 80{,}000$ steps.** Validation Pearson $r$ on human fold 4 (rescaled output space), measured at the closest evaluation step $\leq 80{,}000$. See Figure 3 for the full learning curves and for the caveats concerning the earlier dataset and different training recipe used for these two runs.

| Track | Pretrained | From scratch | $\Delta$ |
|---|---|---|---|
| ATAC | 0.6683 | 0.6521 | +0.0161 |
| CAGE | 0.5334 | 0.5221 | +0.0113 |
| ChIP-histone | 0.7563 | 0.7348 | +0.0215 |
| ChIP-TF | 0.6116 | 0.5948 | +0.0169 |
| DNase | 0.6876 | 0.6621 | +0.0255 |
| PRO-cap | 0.6654 | 0.6507 | +0.0147 |
| RNA-seq | 0.7854 | 0.7563 | +0.0291 |
| **Overall** | **0.6721** | **0.6524** | **+0.0197** |

mapping. This step is necessary because $\log(1+x)$ is not scale invariant, so applying it to two differently scaled versions of the same signal yields different correlations. Splice-site usage is unaffected, being a fraction in $[0, 1]$ in both label sets.

**Agreement with AlphaGenome's released labels.** Because our non-splice related labels were reproduced independently from the primary data sources, we verified them against the labels AlphaGenome subsequently released for the same intervals. Scoring each track by Pearson $r$ across all positions pooled over the 1,576 fold-3 intervals and averaging over tracks, the two label sets agree to $r = 1.000$ for ATAC, PRO-cap and all 2,733 ChIP-seq tracks, $r = 0.9997$ for RNA-seq and $r = 0.998$ for DNase. CAGE agrees slightly less closely ($r = 0.99$, or 0.93 after the $\log(1+x)$ transform), which is expected since CAGE was re-aggregated independently from FANTOM5; the residual difference reflects aggregation choices (which libraries are pooled per cell type, normalization, and strand conventions) rather than a difference in the underlying sequencing data. AlphaGenome did not release its processed GTEx-derived RNA-seq tracks, so those 54 tracks are excluded both from this comparison and from every reported RNA-seq score, which therefore covers the 613 ENCODE-derived RNA-seq tracks of the 667 we train on. The licensing terms are the same for both label sets, since both are processed from the same public sources; consistent with the GTEx General Research Use terms, no GTEx-derived ground-truth data are included in either release.

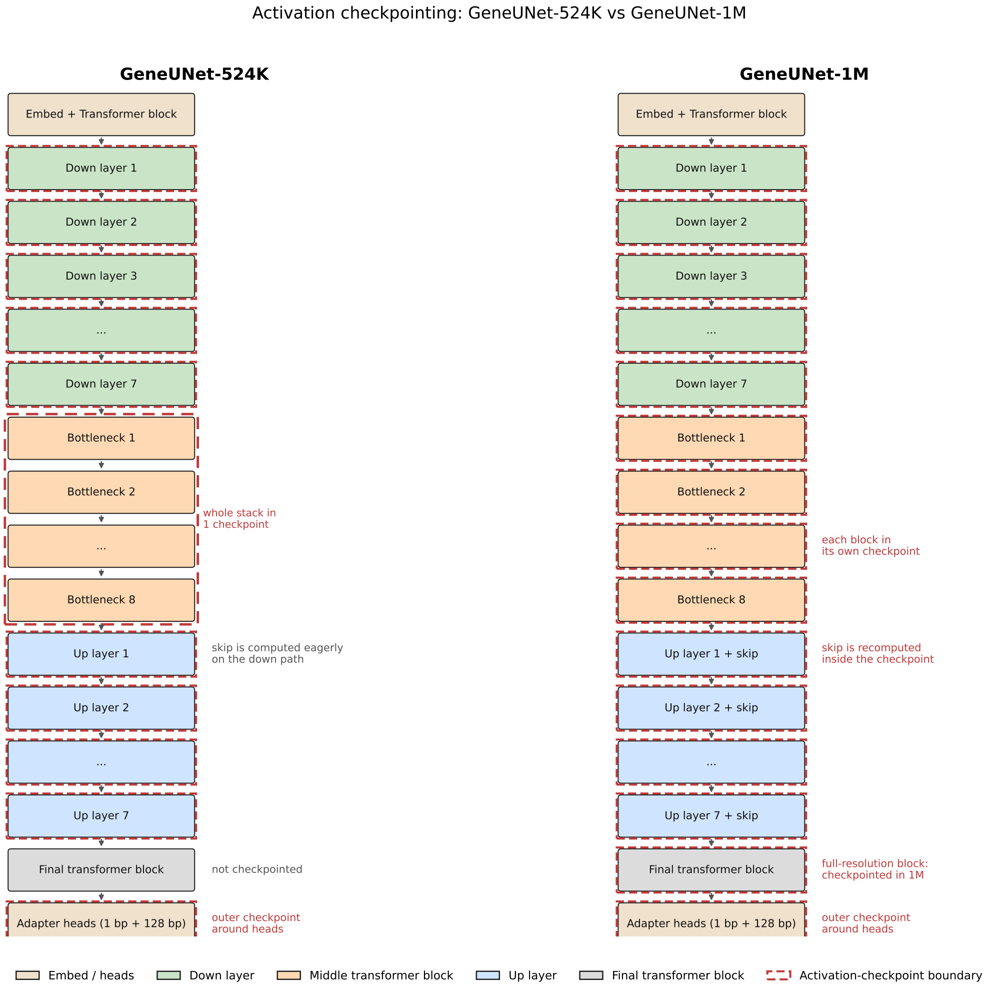


Figure 2 | Activation checkpointing strategy for the 524 kb phase (left) and the 1 Mb phase (right). Dashed red outlines indicate a single activation-checkpoint boundary: activations inside the outline are freed on the forward pass and rematerialised on backward. The 524 kb phase wraps the entire middle transformer stack in a single checkpoint and computes U-Net horizontal skip connections eagerly on the down path. The 1 Mb phase uses per-block middle checkpointing, defers horizontal skips to a combined up+skip checkpoint, and checkpoints the final transformer block — four changes needed to fit the 2× longer activations on a single 8-GPU H100 node.

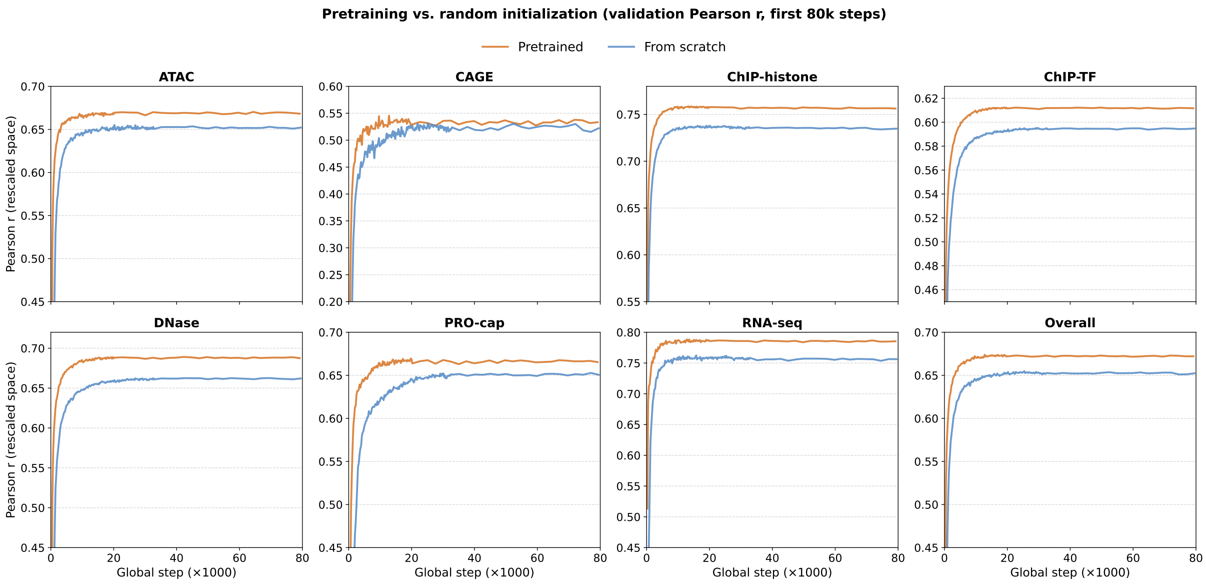


Figure 3 | **Effect of multi-species pretraining on AlphaGenome fine-tuning.** Validation Pearson $r$ (in the model's rescaled output space) on a subsample of human fold 4 over the first $80{,}000$ optimizer steps of two otherwise-identical 524 kb fine-tuning runs: one starting from the OpenGenome2 pretrained checkpoint (orange) and one from random initialization (blue). Per-track panels show the seven AlphaGenome track categories; the *Overall* panel aggregates across tracks. The pretrained run maintains a consistent advantage across all tracks throughout training. These runs used an earlier version of the training data and a different recipe than the main-text runs, so absolute values are not directly comparable to the main fold-3 test numbers; the comparison between the two curves is, however, internally consistent.
